# Combining epiGBS markers with long read transcriptome sequencing to assess differentiation associated with habitat in *Reynoutria* (aka *Fallopia*)

**DOI:** 10.1101/2020.09.30.317966

**Authors:** Marta Robertson, Mariano Alvarez, Thomas van Gurp, Cornelis A. M. Wagemaker, Fahong Yu, David Moraga Amador, William G. Farmerie, Koen J. F. Verhoeven, Christina L. Richards

## Abstract

Despite the limitations of genetic bottlenecks, several invasive species appear to thrive in non-native ranges with extremely low levels of sequence-based genetic variation. We previously demonstrated differentiation of DNA methylation to habitat types of the highly clonal, genetically depauperate Japanese knotweeds using anonymous markers, but the functional relevance of this DNA methylation variation is unknown. Here, we sequenced the full transcriptome combined with a reduced representation bisulfite sequencing approach, epigenotyping by sequencing (epiGBS), to characterize the association among DNA methylation, functional transcripts and the diverse habitat types occupied by the invasive *Reynoutria* species. We identified 50,435 putative transcripts overall, of which 48,866 were annotated with the NCBI NR database. Of these 17,872 (35%) and 16,122 (32%) transcripts shared sequence identity with *Arabidopsis thaliana* and *Beta vulgaris*, respectively. We found genetic differentiation by habitat type suggesting the action of selection and a marginal pattern of differentiation of DNA methylation among habitats, which appears to be associated with sequence differences. However, we found no individual methylation loci associated with habitat, limiting our ability to make functional interpretations. Regardless of the source of variation in DNA methylation, these changes may represent an important component of the response to environmental conditions, particularly in highly clonal plants, but more fine scale genomics analysis is required to test if DNA methylation variation in this system is responsible for functional divergence.

## Introduction

Following introduction outside their native range, invasive species populations are often severely reduced in genetic variation compared to their native populations, thus theoretically decreasing their evolutionary potential. However, paradoxically, these species are often successful colonizers of new habitats. High migration rates, repeated introductions, “general purpose” or “pre-adapted" genotypes, hybridization with native species, phenotypic plasticity, and uniparental reproductive modes (self-fertilization and asexual propagation) are among the factors identified in different systems that have been proposed to resolve this paradox (Richards *et al.* 2006; van Kleunen *et al*. 2010, 2018; Gurevitch *et al*. 2011; Barrett 2015; Estoup *et al*. 2016). Understanding the complexities of invasive species, and the multifaceted genomic mechanisms that contribute to invasion success has become a vibrant area of research in evolutionary ecology and environmental sciences.

Ecological genomics studies have described the distribution of genetic diversity for many species, detecting genomic variants that contribute to local adaptation and phenotypic variation (Feder & Mitchell-Olds 2003; Andrew *et al*. 2013; e.g. Roda *et al*. 2013; Colautti and Lau 2015). However, despite broad patterns of genetic diversity on the landscape, biologists have only a limited understanding of the actual molecular underpinnings of organismal responses to complex biotic and abiotic factors (Pigliucci 2010; Keller 2014). In an effort to answer these questions, ecologists have applied transcriptomic approaches to examine gene expression in natural environments (Alvarez *et al.* 2015; Ferreira *et al*. 2017) shedding light on the expression differences that underlie divergence and adaptation (e.g. Lai *et al.* 2006; Elmer & Meyer 2011; Andrew *et al.* 2013). Therefore, ecological transcriptomics studies have moved one step closer to understanding the molecular mechanisms underlying adaptive traits, but the regulation of transcription in complex genomes largely remains a mystery.

Changes in gene expression can mediate or be mediated by epigenetic changes (chromatin modifications, DNA methylation, small RNAs), which can vary among individuals within populations. These epigenetic changes are sometimes heritable (Schmid *et al.* 2018); and, even when they are not inherited with fidelity, can be correlated with other transient yet important phenotypic outcomes (Wibowo *et al.* 2016). DNA methylation increases in variance in response to allopolyploidy (Salmon *et al*. 2005), and exposure to stress (Verhoeven *et al*. 2010), and may effect ecologically important phenotypes (Johannes *et al.* 2009; Cortijo *et al.* 2014; Richards *et al.* 2017). Variation in DNA methylation has also been correlated to habitat types and shifts in species range (Xie *et al*. 2015; Foust *et al*. 2016; Jueterbock *et al.* 2020). As both a regulator of gene expression and a modification resulting from gene expression (Aceituno *et al.* 2008, Zhang *et al.* 2018), environmentally induced changes in DNA methylation may be indicative of phenotypic plasticity, which can provide a source of variation when there is little genetic variation (Richards *et al*. 2012; Liebl *et al*. 2013; Jueterbock *et al.* 2020). Phenotypic plasticity may be particularly important for rapid response of invasive species to novel habitats (Richards *et al*. 2006, Nicotra *et al.* 2010; Richards *et al*. 2017). However, little is known about how variation in DNA methylation may translate into function in real environments partly because there is limited information about most plant genomes, and the responses that are important for tolerance of biotic and abiotic factors are often multi-genic (Baxter & Dilkes 2012; Assmann 2013; Alvarez *et al*. 2015).

In previous work, we measured the phenotypic and genotypic variation from several populations of invasive Japanese knotweed (*Reynoutria japonica* and *R. x bohemica*). We discovered phenotypic differences and epigenetic divergence of clonal individuals with the same genotype from different habitats (Richards *et al.* 2008, 2012). Overall, our findings were consistent with epigenetically based differentiation in response to habitat, however, several alternative explanations exist for why plants from different habitats can show different DNA methylation profiles. For example, epigenetic mutations could accumulate stochastically among different populations or epigenetic differences might reflect differences in gene expression among different habitats. Epigenetic differences might also reflect genetic differences among habitats that were undetected by our AFLP study; clonal individuals might accumulate genetic differences through somatic mutations which could impact methylation patterns, but our ability to detect changes in DNA sequence was limited by the number of AFLP markers (~200 polymorphic loci). Additionally, SNPs present in one or more sub genomes may not be detectable via AFLP markers since they are dominant markers (Dufresne *et al.* 2014). Further, although our previous study identified population-level patterns of methylation divergence, we were unable to associate the anonymous MS-AFLP loci with gene function (Schrey *et al.* 2013; Paun *et al*. 2019). In this study, we further scrutinize the methylation differentiation among habitats in these populations of knotweed and attempt to elucidate the basis for these differences.

Here, we used a reduced representation bisulfite sequencing technique, epigenotyping by sequencing (epiGBS; van Gurp *et al.* 2016), to measure genetic and DNA methylation differentiation among individuals previously identified as clonal replicates, and developed a transcriptome using the PacBio long read sequencing technology. In order to investigate methylation changes that correlated to habitat, we focused on individuals representing a single genotype of *R. japonica* and individuals representing a single genotype of *R. x bohemica* based on previous AFLP analysis (Richards *et al.* 2012). With this technique and increased resolution, both in number and in detail of the markers, we sought to investigate more accurately the genome-wide DNA methylation changes within each clone, the changes associated with habitat, and if these changes could be associated with candidate genes to indicate potential for regulation of gene expression. We also examined the possibility of DNA sequence differences within AFLP defined clones that were not apparent in our previous study, that might be associated with habitat and DNA methylation.

## Materials and Methods

### Reynoutria species complex

Historically, the taxonomy of the Japanese knotweeds has been complicated (see Richards *et al*. 2008; Schuster *et al.* 2011). *Reynoutria japonica* Sieb. & Zucc. aka *Fallopia japonica* (Houtt.) Ronse Decraene (2N=44 or 88) is considered a primary colonizer, important in the establishment of vegetation on newly formed, bare volcanic habitat. Japanese populations of *R. japonica* are extremely variable in morphology and at the molecular level (Bailey 2003). The distribution of the closely related species *R. sachalinensis* (2N=44) is restricted to the Sakhalin Islands (Russia) and northern Japan, typically found along fresh-water waterways (Bailey 2003). Hybrids (*R. × bohemica* Chrtek & Chrtková; 2N=44, 66, or 88), are much more common in the invasive range of Europe, with greater genetic diversity and more rapid spread than either parent in the invasive range (Mandák *et al*. 2005). In the U.S., recent studies suggest that spread of all three taxa takes place through both vegetative and sexual reproduction and that the morphological variation is much greater than that reported in Europe (Gammon *et al*. 2007; Grimsby *et al*. 2007; Gammon & Kesseli 2010).

In a previous study, we collected Japanese knotweed s.l. from five marsh sites, five beach sites and three roadside sites across eastern NY (Suffolk County, Long Island and Westchester County), Connecticut, and Rhode Island (Table 1). Because methylation is environmentally labile, we collected rhizomes from the field and grew fresh leaf tissue in the greenhouse. At each site, we collected rhizomes approximately 10 m apart to maximize the likelihood of collecting different genotypes, and to represent the full area of each of the 13 sites. In the subsequent AFLP analyses, we identified that all *R. japonica* that we sampled belonged to one haplotype (haplotype F) which was found in all three habitats. In addition, one haplotype (G) of *R. x bohemica* was abundant in several sites across all three habitats (Richards *et al*. 2012). We also surveyed two to four individuals from each of the populations to confirm species or hybrid classes based on cytology and morphology (Richards *et al.* 2008; Richards *et al*. 2012).

**Table 1.** Sampling information across all sites.

| Site | Location | Latitude | Longitude | N | Species | Transcriptome<br>(tissues) |
| --- | --- | --- | --- | --- | --- | --- |
| **Beach**<br>**(16)** |  |  |  |  |  |  |
| HBCT | Madison, CT | 41 16.17 | 72 33.39 | 3 | *F. japonica* | None |
| HP | Southold, NY | 41 05.17 | 72 26.68 | 3 | *F. japonica* | Flower |
| MSH | Port Jefferson, NY | 40 57.74 | 73 02.58 | 2 | *F. × bohemica* | Flower, rhizome |
| PJB | Port Jefferson, NY | 40 57.88 | 73 03.17 | 6 | *F. japonica*/<br>*F. × bohemica* | Leaf |
| RPB | Rocky Point, NY | 40 57.96 | 72 57.28 | 2 | *F. × bohemica* | Flower |
| **Marsh**<br>**(19)** |  |  |  |  |  |  |
| CBH | Port Jefferson, NY | 40 57.19 | 73 02.71 | 7 | *F. japonica*/<br>*F. × bohemica* | Leaf |
| CMM | Center Moriches,<br>NY | 40 47.96 | 72 46.41 | 3 | *F. japonica* | Flower, leaf |
| OLCT | Lyme, CT | 41 18.69 | 72 20.10 | 1 | *F. japonica* | None |
| RHBH | Riverhead, NY | 40 54.24 | 72 37.11 | 4 | *F. × bohemica* | Leaf, rhizome |
| WH | Brookhaven, NY | 40 46.23 | 72 53.86 | 4 | *F. × bohemica* | None |
| **Roadside**<br>**(12)** |  |  |  |  |  |  |
| CMR | Center Moriches,<br>NY | 40 47.98 | 72 46.42 | 4 | *F. japonica* | Leaf, rhizome |
| HL | Rocky Point, NY | 40 57.82 | 72 57.29 | 0 | *F. × bohemica* | Leaf |
| RHC | Riverhead, NY | 40 54.53 | 72 37.47 | 4 | *F. × bohemica* | Leaf, rhizome |
| ST | Smithtown, NY | 40 51.47 | 73 12.60 | 4 | *F. × bohemica* | Rhizome |

### Iso-Seq Transcriptome

In order to maximize the representation of different types of transcripts in the transcriptome, we collected three different tissues (rhizome, leaf and flower) from 11 natural populations of plants growing under different field conditions over the span of three days in September 2014. We extracted RNA with the Spectrum Plant Total RNA kit (Sigma-Aldrich) according the manufacturer’s protocol. In total, we used 17 samples from flower (4), leaf (7), and rhizome (6) tissues collected across 11 sites in beach, roadside and marsh habitat that met quality standards (Table 1). We pooled equal masses from each sample into one single tube for Iso-Seq processing.

We constructed full-length, RNA sequencing libraries using Iso-SeqTM (Pacific BioSciences) according to the recommended protocol by PacBio (Tilgner *et al.* 2013, Schreiner *et al.* 2014), with a few modifications. Briefly, we used only RNA preparations with an RNA Integrity Number (RIN) >= 3.7, as indicated by the Agilent BioAnalyzer or TapeStation. If needed, we used the ZYMO Research RNA Clean and Concentrator kit (Cat. # R1015) to concentrate samples that were too dilute. Pure RNA preparations that were considered suitable for IsoSeq typically displayed absorbance ratios of A260/280= 1.8-2.0 and A260/230> 2.5. We synthesized full-length cDNA using the Clontech SMARTer PCR cDNA Synthesis kit (Cat. # 634925). When starting with one microgram of pure/intact RNA, approximately 15-18 PCR cycles were required to generate 10-15 micrograms of ds-cDNA. We size-selected total cDNA on the SageELF instrument (SageScience), using the 0.75% Agarose (Native) Gel Cassettes v2 (Cat# ELD7510), specified for 0.8-18 kb fragments. Fragments were collected in four size ranges: 0.7-2 kb, 2-3 kb, 3-5 kb, and >5kb. Initial cDNA amplification generated 0.5-1.0 microgram for the two smaller size bins, a sufficient amount for SMRT bell sequencing library construction. Fragments from the larger size bins (2-3 kb and >5kb) were submitted to additional amplification cycles (12 and 14 cycles, respectively). This second amplification resulted in approximately 1.5 microgram of cDNA for the 3-5 kb fraction, and no detectable amounts for the 5-10 kb fraction. Independent SMRT bell libraries were constructed for each one of the 0.7-2 kb, 2-3 kb and 3-5 kb cDNA fractions. The library from each of the three fractions (0.7-2 kb, 2-3 kb, 3-5 kb) was sequenced separately on 2, 3, and 3 SMRT cells (v3) respectively (PacBio RSII, DNA polymerase binding kit p6 v2, DNA sequencing reagent 4.0 v2).

We cleaned samples of SageELF fractions and throughout SMRT bell library construction using AMPure magnetic beads (0.6:1.0 beads to sample ratio) and eluted final libraries 15 ul of 10 nM Tris HCl, pH 8.0. Library fragment size was estimated by the Agilent TapeStation (genomic DNA tapes), and this data was used for calculating molar concentrations. Between 100-200 pM of library was loaded onto the PacBio RS II sample plate for sequencing. All other steps in for sequencing were done according to the recommended protocol by the PacBio sequencing calculator and the RS Remote Online Help system. Each SMRT cell generated 75-90 thousand polymerase reads.

We processed the raw reads generated from multiple insert-size libraries by PacBio RSII sequencer with the PacBio SMRT portal system (version 6.0). The reads of inserts from subreads, including the full-length non-chimeric reads, were produced by RS_IsoSeq (Minoche *et al.* 2015). We applied the iterative clustering for error correction (ICE) algorithm and Quiver for improving isoform accuracy and removing redundancy. We further processed the sequences of both the full-length and fragments of high-quality isoforms with PTA version 3.0.0 (Paracel Transcript Assembler; Paracel Inc, Pasadena, CA).

In PTA, the low-quality bases were trimmed and the sequences with length <75bp and the plastids and ribosomal RNA genes of plants were excluded from further analysis. The cleanup sequences were clustered and assembled based on the CAP3-based PTA assembly module. We assessed the completeness of the assembled transcriptome with BUSCO v3 (Benchmarking Universal Single Copy Orthologs) using the plant set (Embryophyta) as a database of BUSCO group with 1,440 genes (Seppey *et al.* 2019).

We searched the NCBI NR (blastx) and NT (blastn) databases to annotate the consensus sequences resulting from the PTA with e-value <=1e-4 as a cutoff. We used the best scoring hit from the top 25 hits for each query sequence for GO assignment. These GO term assignments were organized around the GO hierarchies that are divided into biological processes, cellular components, and molecular functions. In addition, we also characterized the assembled sequences with respect to functionally annotated genes by BLAST searching against the NCBI reference sequences (RefSeq) of *Arabidopsis thaliana* (35,375 transcripts) and *Beta vulgaris* (78,200 transcripts).

### EpiGBS library construction and Illumina sequencing

We isolated DNA from 47 samples (19 *R. japonica* and 28 *R. x bohemica*) from among 13 sites using the Qiagen DNeasy plant mini kit according to the manufacturer’s protocol. Following library preparation, we removed five samples due to stochastic undersequencing that lowered the number of available high-quality loci in those individuals, for a final sample size of 42 individuals (Table 1). We prepared epiGBS libraries *sensu* van Gurp *et al.* (2016). Briefly, isolated DNA was fragmented with the enzyme PstI, which is sensitive to CHG methylation and biases resulting libraries toward coding regions (van Gurp *et al.* 2016). After digestion, adapters with variable barcodes were ligated to either end of the resulting fragments. Adapters contained methylated cytosines to ensure their sequence fidelity through the subsequent bisulfite treatment. We used the Zymo EZ Lightning methylation kit to bisulfite treat and clean the DNA. Libraries were then amplified with the KAPA Uracil Hotstart Ready Mix with the following PCR conditions: an initial denaturation step at 98°C for 1 min followed by 16 cycles of 98°C for 15s, 60°C for 30s, and 72°C for 30s, with a final extension of 72°C for 5 min. We used rapid run–mode paired-end sequencing on an Illumina HiSeq2500 sequencer at Wageningen University & Research using the HiSeq v4 reagents and the HiSeq Control software (v2.2.38), which optimizes the sequencing of low-diversity libraries (van Gurp *et al.* 2016). We ran our 47 knotweed samples in a single lane that included a lambda control and 48 samples of another species (*Spartina alterniflora*) prepared with the same protocol for another study (Alvarez, Robertson *et al.* 2020).

### Data pre-processing

We used the pipeline published by van Gurp *et al.* (2016) to demultiplex samples, trim adapter sequences, and assemble the *de novo* reference sequence (legacy scripts archived at https://github.com/AlvarezMF/2020_Knotweed_epiGBS). We mapped our de novo reference epiGBS fragments to our *Reynoutria* transcriptome using BLAST with e-value <=1e-4 as a cutoff (Altschul *et al.* 1997). We tabulated SNP and methylation polymorphisms using the epiGBS analysis pipeline. We removed SNP and methylation loci with less than 10x depth of coverage in each individual. Then, both SNP and methylation data were filtered separately to include only loci that were present in more than 60% of individuals, with no more than 80% missing from any one individual after filtering loci. Thus, each locus had, at most, 40% missing data, while each individual had less than 80% missing data across all loci. The remaining missing data (Fig. S2) were imputed using a k-nearest neighbors approach (Hastie *et al.* 2019). Additionally, we conducted separate analyses for individuals previously identified as haplotype F of *R. japonica* and those identified as haplotype G of *R. x bohemica* (Richards *et al.* 2012). In these analyses, loci were filtered again, separately within subsets, yielding a new set of common SNP and methylation loci. All genome-wide analyses were repeated in these subsets.

There were several sources of uncertainty in genotyping for this non-model plant that has no reference genome even when using allele frequencies: stochastic under-sequencing, the high ploidy of the individuals being sequenced, a mapping approach based on local de novo assemblies generated from bisulfite reads, and a novel pipeline. To assess the robustness of our results, we repeated genome-wide analyses on all samples combined, as described below, with stricter filtering thresholds on 31 of the original 47 samples. After removing SNP and methylation loci with less than 10x depth of coverage in each individual, we filtered both SNP and methylation data separately to include only loci that were present in more than 90% of individuals, with no more than 50% missing from any one individual after filtering loci. Thus, each locus had, at most, 10% missing data, while each individual had less than 50% missing data across all loci. All analyses were highly concordant, with significant effects in the same direction and magnitudes as described below.

### Population genetics

All statistical analyses were performed in R v 3.5.3 (R Core Team, 2017). The epiGBS technique, and the sequencing design that we chose, did not provide sufficient sequencing depth to estimate polyploid genotype likelihoods with confidence, particularly considering the lack of a high-quality reference genome (Boutte *et al.* 2016; Dufresne *et al.* 2014). We therefore used the frequency of the most common allele at each polymorphic locus as a substitute for genotype at each locus, as discussed in Alvarez & Robertson *et al.* (2020).

We obtained pairwise F_ST_ values between populations to test for significant differentiation (StAMPP, Pembleton *et al.* 2013). We also used distance-based redundancy analysis (RDA function in the Vegan package v. 2.5-2; Oksanen *et al.* 2017) to minimize false positives (Meirmans 2015) in assessing isolation by distance using the formula (genetic distance ~ latitude + longitude). We visualized data using principal components analysis (PCA). We also visualized relationships between samples using hierarchical clustering on Euclidean distances and generated heat maps and histograms based on Jaccard similarities between individuals, all implemented in R (R Core Team 2017).

To quantify the relationship between genome-wide variation and habitat, we used partial constrained redundancy analysis (RDA, implemented with the RDA function in the Vegan package v. 2.5-2; Oksanen *et al.* 2017). RDA is a multivariate ordination technique that allowed us to assess the joint influence of all SNPs simultaneously, while effectively controlling for both population structure and false discovery (Forester *et al.* 2018). We attempted to control for variation among sites with a replicated sampling strategy, but rather than using a single term for “population”, we conditioned our ordination on variables identified by latent factor mixed models analysis using the LFMM package (Caye *et al.* 2019), which provides a method to account for residual variation due to unmeasured differences among individuals, including population structure, environmental variation, life history variation, and geographical separation (Leek *et al.* 2017). Latent factors were modeled with K=3 (K=4 for *R.x bohemica* subset), as determined by scree plot. We used RDA to fit our final model with the formula (SNP matrix ~ habitat + Condition(latent factors)). We used a permutational test (9999 permutations; Oksanen *et al.* 2017) to assess the likelihood that individuals in different habitats differed by chance, and assumed that non-random differentiation was consistent with, but not conclusive evidence of, the action of selection in different habitats. We visualized results using principal components analysis and the RDA axes, which represent the lines of maximal differentiation between habitat types. We also tested for the significance of RDA axes via permutation tests, which provide an additional assessment of non-random differentiation between sites. To identify SNPs that were significantly correlated with habitat type, we used LFMM (Caye *et al.* 2019) to identify outlier loci. P-values were adjusted for genomic inflation factor and corrected for multiple testing via Q-value (Storey *et al.* 2015), at a significance threshold of 0.05.

### Methylation analysis

During the filtering process, loci were annotated with their sequence context (CG, CHG or CHH), but all contexts were pooled for distance-based analyses as well as family-wise error rate corrections after locus-by-locus modeling. We prepared methylation data for analysis by first imputing missing data using the same approach as for genetic data. We then quantified methylation frequency, defined as the fraction of methylated cytosines observed out of the total number of cytosines measured at a given locus (methylated cytosines/(methylated+unmethylated cytosines)).

To identify signatures of DNA methylation variation that were correlated with habitat while controlling for genetic structure, we again estimated latent variables with LFMM (Caye *et al.* 2019) as above using K=4 (K=2 for *R. japonica* subset). In addition to the advantages described above, latent factor analysis (or the related surrogate variable analysis) provides a control for cell type heterogeneity in epigenomic studies (Akulenko *et al.* 2016; Caye *et al.* 2019; McGregor *et al.* 2016). We then modeled the impact of habitat on genome-wide patterns of DNA methylation frequencies while controlling for latent variation as well as population structure via RDA (Vegan v. 2.5-2; Oksanen *et al.* 2017) with the formulas (methylation distance ~ habitat type + Condition(latent factors) and (methylation distance ~ habitat type + Condition(latent factors + the first five principal components of the SNP matrix). By removing the influence of genomic population structure on methylation differences in the second formula, we assumed that the resulting differences would be primarily attributable to DNA methylation variation that was independent of genetic differences. We used a permutational test (9999 permutations; Oksanen *et al.* 2017) to assess the likelihood that individuals in different habitats differed by chance, and visualized results using principal components analysis. We again conducted separate analyses for two groups previously described.

Because different methylation contexts may differ in regulation, we assessed the magnitude of differences due to habitat type in context-specific methylation frequency using linear regressions. To identify methylation polymorphisms that were significantly correlated with habitat type, we used LFMM to identify outlier loci. P-values were adjusted for genomic inflation factor and corrected for multiple testing via Q-value (Storey *et al.* 2015), at a significance threshold of 0.05.

## Results

### Transcriptome of Reynoutria species

The Pacbio long-read isoform sequencing based on 8 SMRT cells over three fraction sizes (07-2kb, 2kb-3kb, and 3-5kb) produced a total of 610,797 reads of inserts with an average length of 1,531 bp, including 259,723 full length reads of inserts. The *de novo* transcriptome assembly generated 50,435 genes/transcripts with an N50 of 2,346 bp. The reads length ranged from 300 to over 10,943 bps. The BUSCO analysis revealed that the assembled genes/transcripts covered completely or partially 60% of the orthologues from plants. The homologue search against the NCBI NR database suggested that a total of 48,866 transcripts were annotated with e-value <= 1e-4, and 4,352 genes were assigned at least one GO category (cellular component, molecular function, and biological process) in Gene Ontology. In addition, the BLAST search over NCBI RefSeq database indicated that about 17,872 and 16,122 genes of *Reynoutria* showed homology with the genes of *Arabidopsis thaliana* and *Beta vulgaris*, respectively.

### epiGBS provides a genome-wide scan of DNA methylation

The *de novo* assembly using the epiGBS pipeline (van Gurp, *et al.* 2016) resulted in 7,924 contiguous fragments of 19-202 basepairs for a total length of 1,219,146 bp (Figs. **S1, S2**). Given the large size of *Reynoutria* genomes (2C values up to 6.48 pg; Bailey & Stace 1992), our reduced representation approach assayed approximately 0.0001% of the genome. We note that fragments that were >90% similar were merged, creating the likelihood of merged polyploid homeologs. Our bisulfite conversion rate was 99.61% per position, as estimated from the lambda phage spike-in.

With BLAST, we found 2,473 fragments mapped to 1,932 transcripts in our *de novo* transcriptome. Of these, 338 transcripts were covered by more than one epiGBS fragment. Of the 2,437 transcriptome-mapped epiGBS fragments, 171 of these fragments overlapped with the 5’ end of transcripts, which has been suggested to harbour regulatory variation in DNA methylation (Neiderhuth *et al.* 2016). However, we found no SNP or methylation outlier loci that were associated with habitat, so we could not make functional interpretations based on reads that overlap with the transcriptome in this data set.

Because of stochastic sample dropout during sequencing, we removed 5 individuals from the final analysis, resulting in 42 total samples. We quantified allele frequency at 101,189 SNP loci across both haplotypes, which were filtered to yield 14,718 informative SNP loci (9,134 in *R. japonica* and 11,123 in *R. bohemica* in sub-analyses). Of these, 243 SNPs occurred in transcripts. Simultaneously, after filtering and imputing missing data, we obtained 17,381 usable methylation loci from an original set of 325,332 observed loci (Fig. **S4**). We quantified methylation frequency at 2,740 loci in the CG context, 4,220 loci in the CHG context, and 10,351 loci in the CHH context. Methylation calls were collapsed for symmetric CG and CHG loci across “watson” and “crick” strands so that methylation on either one or both strands was considered as a single locus. In sub-analyses, we quantified 11,309 methylation loci in putative *R. japonica* samples and 13,130 methylation loci in putative *R. bohemica* samples.

### Clonality and genetic differentiation

Combining the two species, we found significant genome-wide differentiation among samples that was correlated to habitat type (Table 2). Habitat explained 3.2% of variance between our samples (Table 2). These results were recapitulated in our PCA visualization, which shows very little separation by habitat along PCA axes 1 and 2, but discernible differentiation along RDA axis 1, which represents the lines of maximal separation between samples due to habitat differences (Figure 3). Pairwise F_ST_ calculations showed that most sites were significantly different from each other (Table 3); however, we found no evidence of isolation by distance (P>0.05 for latitude and longitude). Finally, we found no SNPs that were significantly associated with habitat type using locus-by-locus tests. When we analyzed *R. japonica* and *R. x bohemica* independently, we found significant genome-wide differentiation correlated with habitat type in *R. x bohemica* only, but we found that patterns of isolation by distance were stronger (Table 3).

**Table 2.**
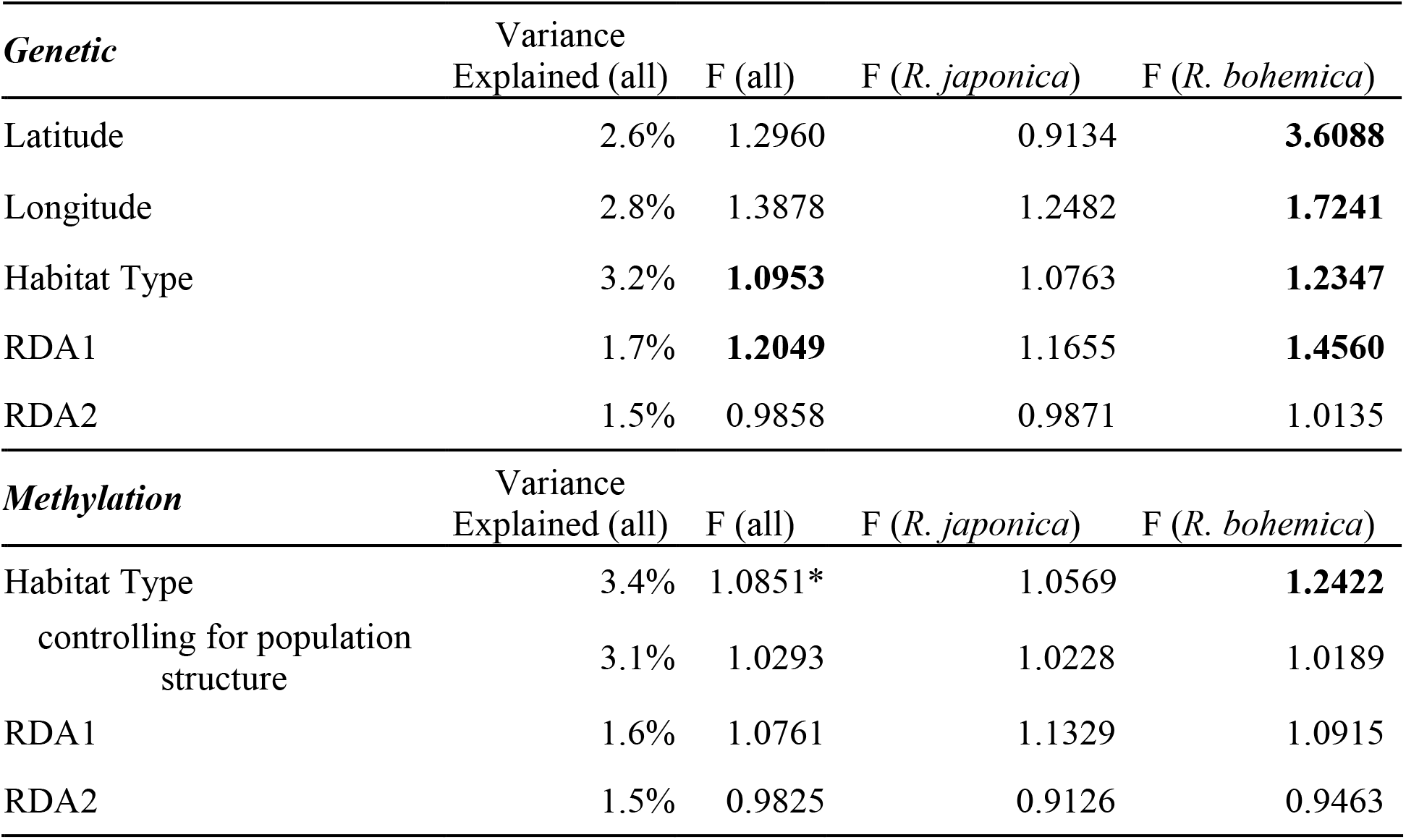
Association between habitat type, genetic distance, and methylation distance for all samples combined. Bolded F values are significant (P < 0.05, *marginal P = 0.057) in permutation tests. Axis tests for methylation data were conducted while controlling for population structure (genetics).

| *Genetic* | Variance<br>Explained (all) | F (all) | F (*R. japonica*) | F (*R. bohemica*) |
| --- | --- | --- | --- | --- |
| Latitude | 2.6% | 1.2960 | 0.9134 | **3.6088** |
| Longitude | 2.8% | 1.3878 | 1.2482 | **1.7241** |
| Habitat Type | 3.2% | **1.0953** | 1.0763 | **1.2347** |
| RDA1 | 1.7% | **1.2049** | 1.1655 | **1.4560** |
| RDA2 | 1.5% | 0.9858 | 0.9871 | 1.0135 |
| *Methylation* | Variance<br>Explained (all) | F (all) | F (*R. japonica*) | F (*R. bohemica*) |
| Habitat Type | 3.4% | 1.0851* | 1.0569 | **1.2422** |
| controlling for population<br>structure | 3.1% | 1.0293 | 1.0228 | 1.0189 |
| RDA1 | 1.6% | 1.0761 | 1.1329 | 1.0915 |
| RDA2 | 1.5% | 0.9825 | 0.9126 | 0.9463 |

**Table 3.** Pairwise Fst among sites. Bold (i.e. all entries) indicates significance at P<0.05.

|  | CBH | CMM | CMR | HBCT | HP | MSH | OLCT | PJB | RHBH | RHC | RPB | ST | WH |
| --- | --- | --- | --- | --- | --- | --- | --- | --- | --- | --- | --- | --- | --- |
| **CBH** |  |  |  |  |  |  |  |  |  |  |  |  |  |
| **CMM** | **0.0428** |  |  |  |  |  |  |  |  |  |  |  |  |
| **CMR** | **0.0247** | **0.0662** |  |  |  |  |  |  |  |  |  |  |  |
| **HBCT** | 0.0004 | **0.0431** | **0.0166** |  |  |  |  |  |  |  |  |  |  |
| **HP** | **0.0405** | **0.0676** | **0.0131** | **0.0216** |  |  |  |  |  |  |  |  |  |
| **MSH** | -0.0072 | **0.0441** | **0.0162** | -0.0099 | **0.0509** |  |  |  |  |  |  |  |  |
| **OLCT** | **0.0292** | **0.0193** | **0.0572** | 0.0057 | **0.0757** | **0.0235** |  |  |  |  |  |  |  |
| **PJB** | **0.0151** | **0.0367** | **0.021** | **0.0246** | **0.0332** | 0.0065 | **0.0451** |  |  |  |  |  |  |
| **RHBH** | **0.0593** | **0.0605** | **0.0291** | **0.0479** | **0.0216** | **0.0721** | **0.0804** | **0.0327** |  |  |  |  |  |
| **RHC** | 0.0009 | **0.0418** | **0.0251** | **0.0119** | **0.0186** | 0.0073 | **0.0484** | **0.0136** | **0.0485** |  |  |  |  |
| **RPB** | **0.0224** | **0.0504** | 0.0186 | 0.0083 | **0.0324** | -0.0032 | **0.0443** | -0.0123 | **0.0247** | -0.0086 |  |  |  |
| **ST** | **0.0345** | **0.0505** | **0.0482** | **0.0285** | **0.035** | **0.0584** | **0.0778** | -0.002 | **0.026** | **0.0267** | -0.007 |  |  |
| **WH** | **0.0176** | **0.0368** | **0.0184** | **0.0196** | **0.025** | **0.0111** | **0.0355** | -0.0035 | 0.0053 | **0.0144** | -0.0283 | 0.0001 |  |

Hierarchical clustering of SNPs showed two major clusters that cleanly separated the previously-identified *R. japonica* and *R. x bohemica* individuals (Figure 2). Within each haplotype, there was also separation by habitat type, but it was less clearly defined (Figures 2, S1). Jaccard similarities ranged from 0.984-0.998 among samples within the same haplotype, which was similar to the range of similarities between haplotypes (0.984-0.994), but also included pairs that were more similar to each other within each haplotype, particularly in *R. japonica* (Figure 1). There were no obvious differences in similarities within versus among sites.

**Figure 1.**
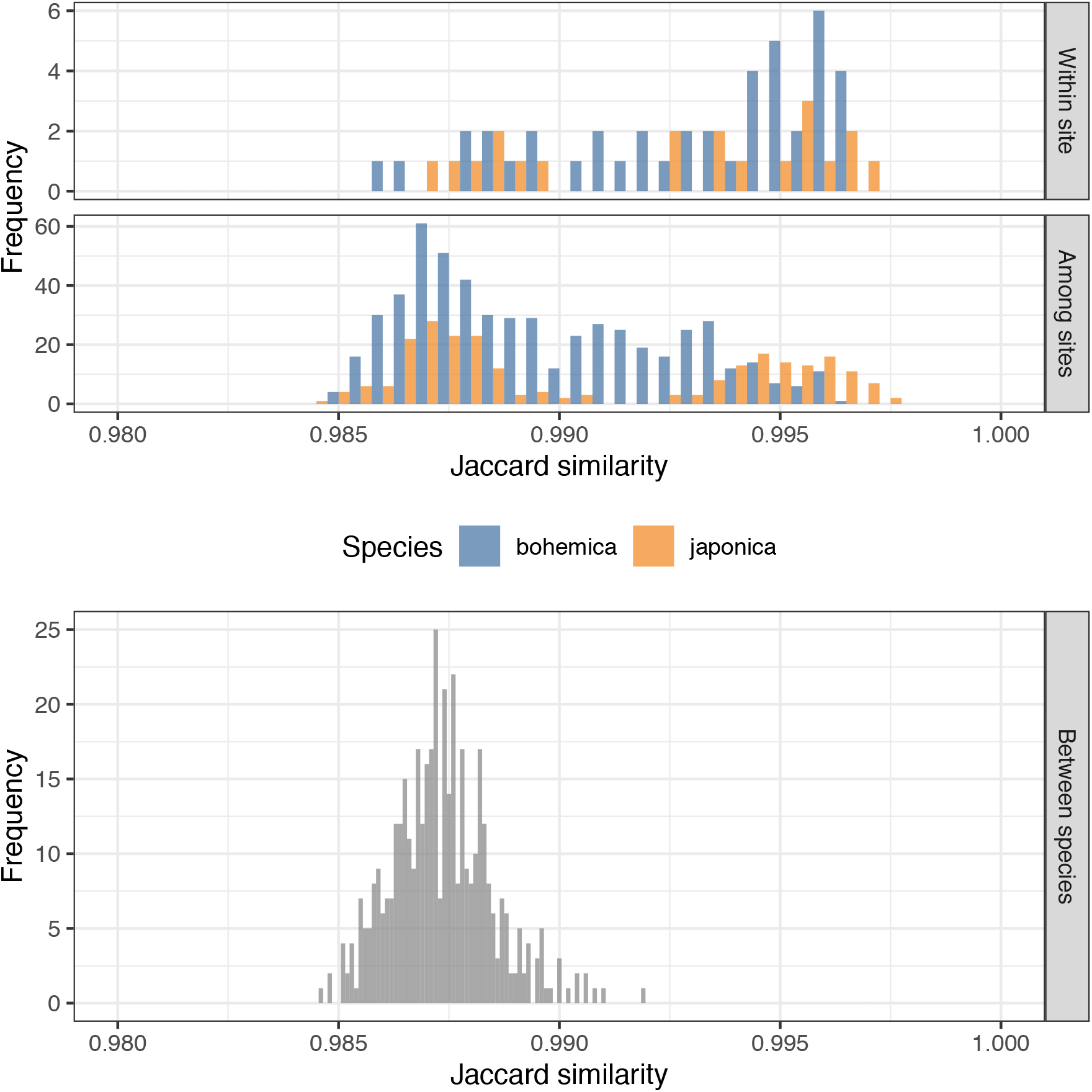
Histogram of Jaccard similarities among individuals within and among sites of each haplotype and between haplotypes of each species previously identified by AFLP (Richards *et al*. 2012).

**Figure 2.**
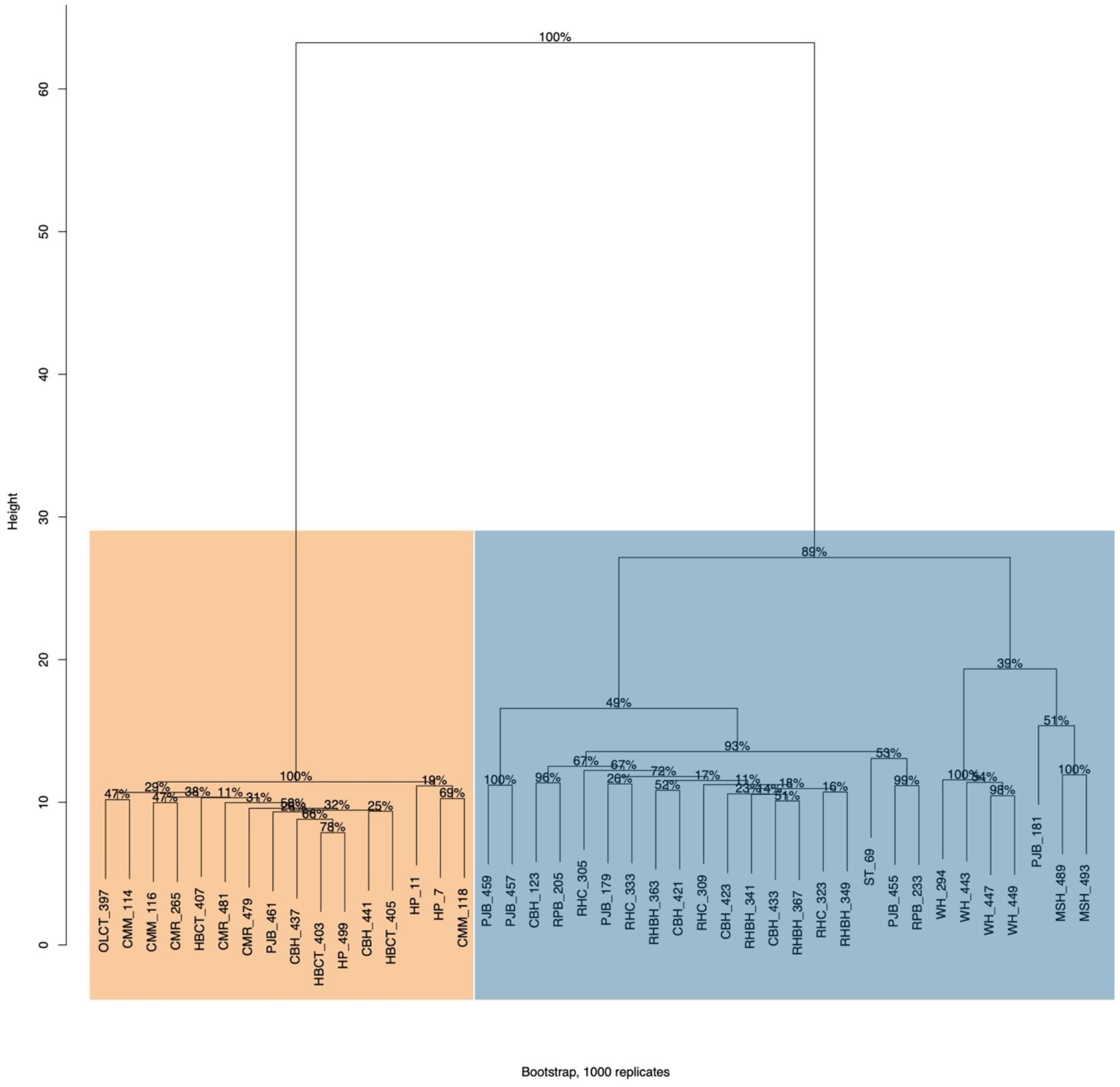
Hierarchical clustering of genetic data. Orange represents individuals previously identified as haplotype F of *R. japonica*, while blue represents individuals previously identified as haplotype G of *R. bohemica* (Richards *et al.* 2012). Percentages represent bootstrap support.

**Figure 3.**
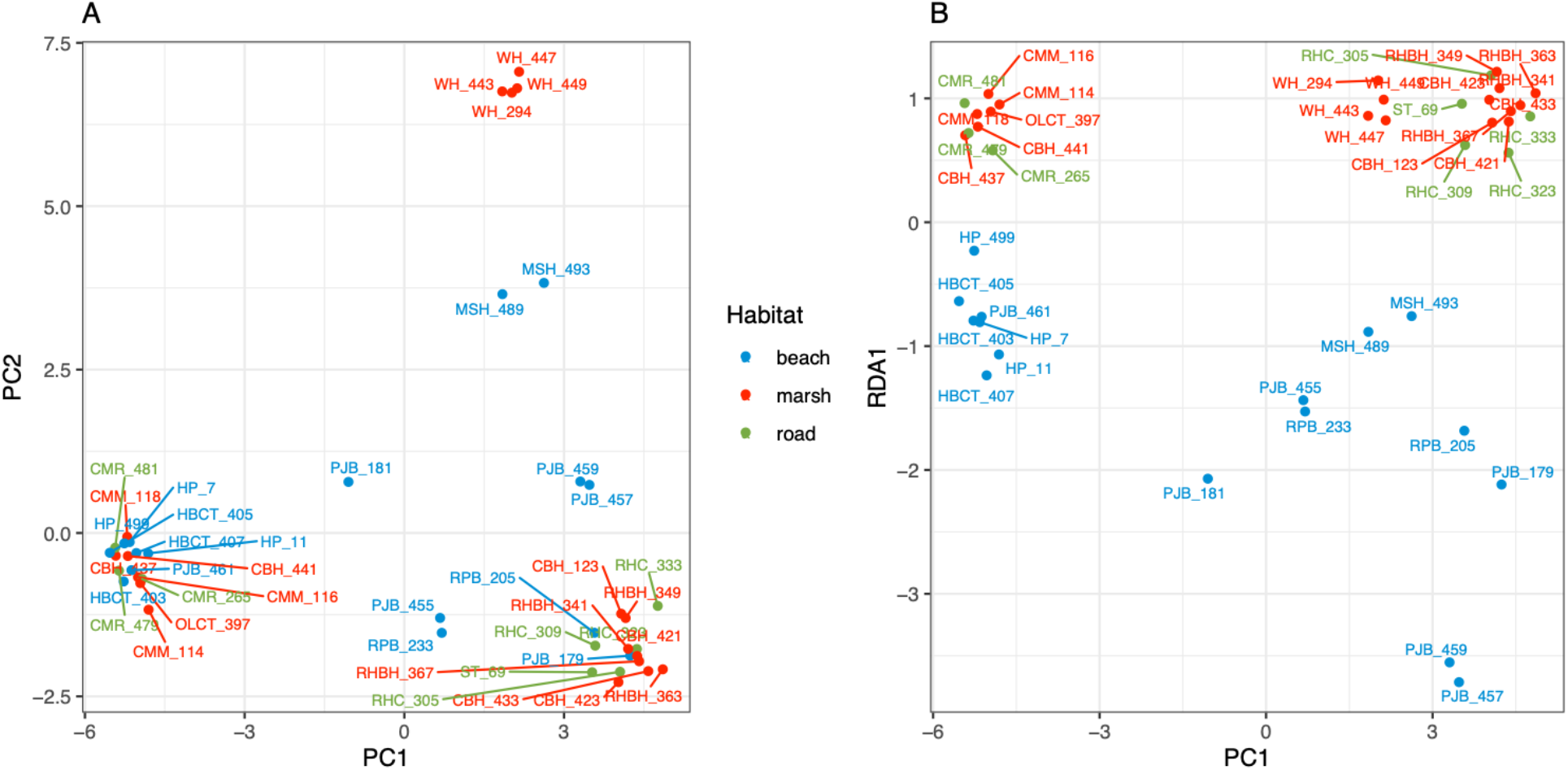
Visualization of genomic differences between individuals by both **A)** Principal components analysis and by **B)** redundancy analysis, where RDA1 represents the line of maximal separation between habitat types.

### DNA methylation differentiation

We found that methylation differences between habitats were marginally significant before controlling for population structure in both overall analyses (P=0.057) and significant (P<0.05) in a separate analysis of samples previously identified as *R. x bohemia*, but we found no significant methylation differentiation by habitat after controlling for population structure (Table 2; Figure 4) both overall and for each species independently. Using locus-by-locus tests for differentiated methylation loci, we found no differentially methylated positions (DMPs, Q<0.05) between habitat types.

**Figure 4.**
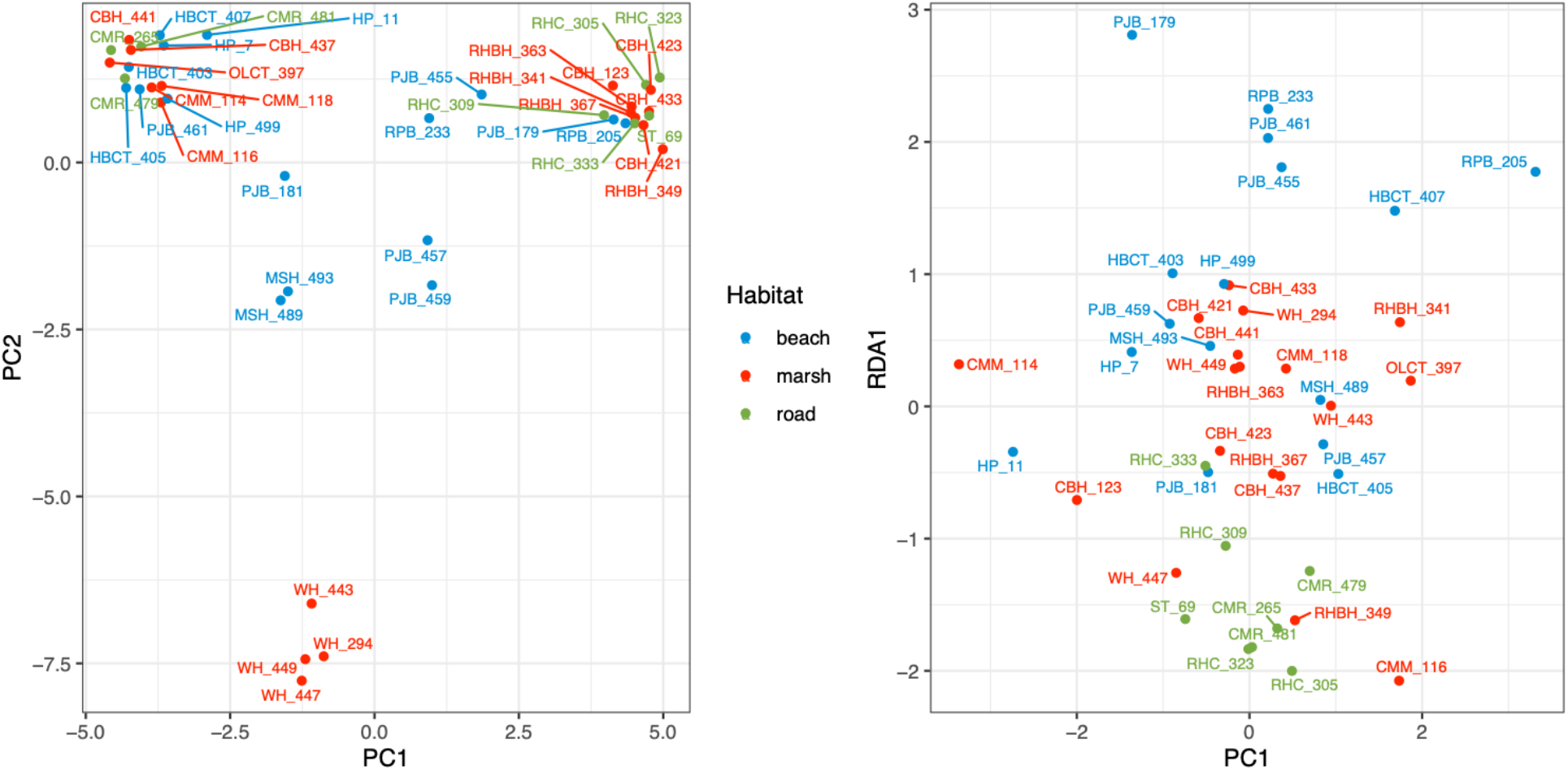
Visualization of epigenomic differences between individuals by both **A)** Principal components analysis and by **B)** redundancy analysis, where RDA1 represents the line of maximal separation between habitat types.

**Figure 5.**
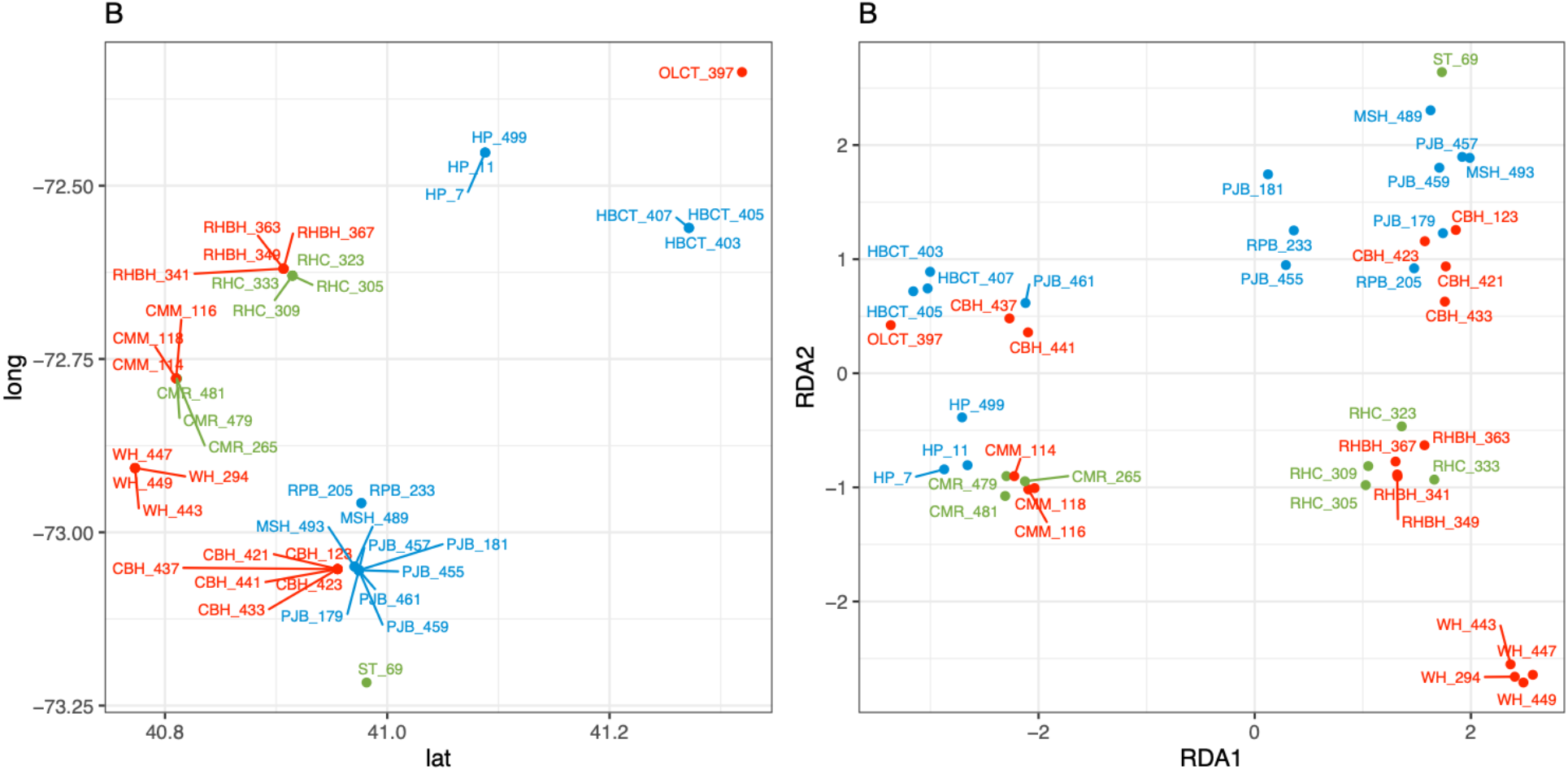
Comparison of patterns of isolation by distance among individuals, as shown by **A)** latitude and longitude, with patterns of habitat differentiation, as shown by **B)** redundancy analysis axes 1 and 2.

## Discussion

Understanding the role of DNA methylation in adaptive divergence is a challenging task. Depending on genetic and molecular context, DNA methylation can be causal to gene expression modulation (Lang *et al.* 2017; Zhang *et al.* 2006), a passive consequence of gene expression (Meng *et al.* 2016, Niederhuth and Schmitz 2017, Secco *et al.* 2015), or without known function. DNA methylation variation can be under genetic control, induced by environmental effects, or can arise spontaneously as epimutations (Johannes & Schmitz 2019; Zheng *et al.* 2017). Because of its ability in some contexts to regulate gene expression, and therefore phenotype, DNA methylation has been proposed to contribute to phenotypic variation, particularly in genetically depauperate populations (Verhoeven and Preite 2014; Douhovnikoff and Dodd 2015; Richards *et al.* 2017). Several invasive species have little genetic variation compared to their source populations, but still have high amounts of phenotypic variation and can colonize diverse habitats (Richards *et al.* 2012; Dlugosch & Parker 2008b). Previous work suggests that epigenetic variation might be induced by environmental stressors, and could contribute to rapidly evolving populations (Verhoeven *et al*. 2010; Richards *et al*. 2012, 2017).

Our previous work in Japanese knotweed showed that despite a lack of genetic variation in invasive populations, the species had successfully colonized diverse habitats, showed a high degree of phenotypic plasticity, and showed an epigenetic signature correlated with source habitat (Richards *et al.* 2008, 2012). However, the genetic and epigenetic variation we examined previously was evaluated via dominant anonymous markers (MS-ALFP), and we sought to more closely examine the identity and potential function of the epigenetic variation underlying the differentiation we detected. In order to do so, we built a transcriptome with the intention of relating the results of our bisulfite sequencing to functional transcripts. Our transcriptome represented approximately 60% of the orthologues from plants according to the software Benchmarking Universal Single Copy Orthologs (BUSCO; Seppey *et al.* 2019), which improved annotation over the shared sequence identity with *Arabidopsis thaliana* (35%) or *Beta vulgaris* (32%) alone. Our 7,924 epiGBS fragments overlapped with 1,932 of these transcripts, suggesting we could find association with function with these libraries. However, we did not identify any associations of SNPs or DMPs with habitat, and we were therefore unable to make functional conclusions about the association of genetic or epigenetic variants with habitat.

In order to isolate epigenetic differences, we sampled one AFLP haplotype of the clonal *R. japonica* and one AFLP haplotype of the clonal *R. x bohemica* from co-occurring habitats and found that they were differentiated genetically from each other. In contrast to our previous work and that of several others that suggested that all invasive populations of *R. japonica* were made up of the same single clone and that *R. bohemica* had very few genotypes (Hollingsworth & Bailey 2000; Mandák *et al.* 2005; Richards *et al.* 2012; Zhang *et al*. 2017), we found genetic differences among individuals within and among populations of both haplotypes. However, we interpret these results with some caution for several technical reasons. First, we note that SNP calls in our species are error prone due to the bisulfite treatment of the reads. Genetic polymorphisms may be confounded with methylation polymorphisms during tabulation, leading to false increases in SNP variation (Gao *et al.* 2015, Liu *et al.* 2012). Second, errors in the *de novo* construction of our reference genome could lead to the appearance of SNP variation where there is none. In absence of an existing reference genome, generating good local reference assemblies based on bisulfite reads in hexaploid and octoploid species presents a significant technical challenge; inaccuracies in the de novo assemblies can generate false positive SNPs in the data that contribute to the impression of genetic divergence between lineages.

Conceptually, our use of allele frequencies rather than genotypes provides some hedge against the use of poor genotyping calls when assessing genome-wide variation, and our discovery of differentiation between populations appears to be robust. However, our estimates of within-clone genetic variation in this study are discordant with several previous molecular marker studies in these species, and we were unable to compare replicate samples of the *same* individuals to identify sources of *within individual* variation in these marker profiles (eg. replicates of the same ramet sensu Douhovnikoff & Dodd 2003). This is important information because genetic diversity that we detected across the assayed individuals could result from polymorphisms within the hexaploid and octoploid genomes that would be difficult to differentiate from differences among multi-locus lineages in our protocol. Still, it is possible that multiple unique clones were involved during multiple *Reynoutria* introduction events, or that there has been substantial somatic mutation since their introduction to the area (Prentis *et al.* 2008; Te Beest *et al.* 2012). These differences may have been missed in our previous study using AFLP markers due to the poor resolution and the dominant nature of those markers (Dufresne *et al.* 2014), and are difficult to address with the current study. However, differentiation between sites suggests that selection is acting on a subset of genetic variation (though the source is unknown). Another recent study also found multiple clonal lineages within other invasive populations of *Reynoutria spp.* (VanWallandael *et al.* 2020). Future studies should include replicate sequencing of individuals known to be clonal replicates *a priori* to better assess within-individual variation in these high ploidy (6x and 8x) *Reynoutria spp.*

Although somatic mutation has traditionally been considered disadvantageous and the somatic mutation rate should be low, recent work suggests that particularly in clonal plants, somatic mutations might be beneficial (Eckert 2002; Fischer & van Kleunen 2002), boosting genotypic diversity and increasing the possibility of beneficial variation (Schoen & Schulz 2019). High genotypic diversity in the clonal plants *Grevillea rhizomatosa* (Gross *et al.* 2012), *Hieracium pilosella* (Houliston & Chapman 2004), and *Zostera marina* (Jueterbock *et al*. 2020; Yu *et al.* 2020) have been attributed to high somatic mutation rates. Although we were not able to measure somatic mutation rate directly in this study, we suspect that the genotypic diversity in invasive Japanese knotweeds could be explained by similar mechanisms. Finer-scale sequencing, combined with a better understanding of the history of introductions in the northeastern US, will be needed to determine the evolutionary history of the species and the ongoing invasion history.

We expected that individuals would be genetically identical within haplotypes, and therefore anticipated little genetic divergence, and only epigenetic differentiation, between sites. However, we found that when evaluating all samples across both haplotypes, beach sites were genetically differentiated from marsh and roadside sites, even after controlling for population structure. This pattern of environmental differentiation was particularly strong in *R. x bohemica*. Because we did not observe similar patterns of isolation by distance across all samples, we suggest that differentiation between habitat types is evidence of, although not conclusive of, the action of selection. When we evaluated each species separately, the effect of habitat was substantially weaker than the effect of latitude in *R. x bohemica*. However, in *R. x bohemica*, the pattern of IBD could be confounded with habitat differentiation, since the habitat types are not dispersed evenly throughout the sampling area. Similarly, *Phragmites australis* has invaded several distinct habitats, and genetic variation is correlated with introduction history as well as habitat type (Guo *et al.* 2018).

Visualization of our redundancy analysis suggests that beach sites are genetically distinct from the other habitat types for both species, but when evaluated in separate analyses, *R. japonica* showed no correlation between genetic variation and environment, suggesting that differentiation is much weaker in *R. japonica* than *R. x bohemica.* The lack of genetic and habitat correlation in *R. japonica* may be due to either weaker selective forces, less effective selection on the lower genetic diversity in *R. japonica*, or low resolution due to the reduced number of samples in the within-species comparison. Conversely, because *R. x bohemica* is a hybrid species, it may harbour more variation that can be subject to selection (Hegarty *et al.* 2006, Prentis *et al.* 2008, Walls 2009). Most notably, these results suggest that despite previous findings that there was no genetic variation in *R. japonica*, the species has been experiencing some divergence in the United States even over the limited time since introductions.

### Epigenetic differentiation

In contrast to our previous study, we did not find an epigenetic signature correlated with habitat that was independent of genetic variation. Instead, our results suggest that the epigenetic differences we did see were correlated with genetic differences present in the population. These results are consistent with another reduced-representation assessment in native populations of *Spartina alterniflora* exposed to the *Deepwater Horizon* oil spill (Alvarez & Robertson *et al.* 2020). We suggest that a less significant epigenetic signature comes from a better characterization of genetic variation and population structure, which is, in turn, enabled by the large increase in the number of genome-wide markers.

In this study, the strong correlation of genotype and epigenotype suggests one of two possibilities. In the first, epigenotype is dependent on genotype (as suggested in Dubin *et al.* 2015, Meng *et al.* 2016; Sasaki *et al*. 2019), and could be reflective of genome-wide effects induced by the introduction to a novel habitat (Colautti & Lau 2015; Dlugosch *et al*. 2015; Estoup *et al*. 2016; van Kleunen *et al*. 2018). Whole genome studies in *Arabidopsis thaliana* and in human cancers have shown that epigenetic variation can be dramatically shaped by single nucleotide changes (Becker *et al.* 2011; Timp and Feinberg 2013; Dubin *et al.* 2015; Feinberg *et al.* 2016; Sasaki *et al.* 2019). In clonal plants, the low level of genetic variation that arises from somatic mutations in natural clonal lineages cannot be excluded since several studies have reported that high rates of somatic mutation may allow asexual species to maintain abundant genetic variation and adapt to changing environmental conditions (Lynch *et al.* 1984; Gill *et al.* 1995; Schoen and Schulz 2019). Genetic control of epigenotype could originate either through nucleotide variation in key epigenetic genes, such as the RdDM pathway (Matzke & Mosher 2014), or through simple changes in cytosine content, which alters the number of possible positions for methylation changes to occur. Joint genetic and epigenetic processes, such as transposable element release or chromatin rearrangements, could be responsible for the correlation between genetic and epigenetic variation (Panda *et al.* 2016), but more fine-scale experiments using alternative techniques would be required to address this hypothesis.

In the second possibility, clonal lineages accumulate stable changes in methylation loci (“epimutations”) in a way that is similar to the random accumulation of sequence mutations. Two samples that are more similar genetically will also be more similar epigenetically but not because of a causal link between genetic variation and methylation variation. In that case, we would find that epigenetic variation is explained by genetic variation in a statistical sense, but that does not necessarily mean that it is determined in a mechanistic sense. Much of our understanding of spontaneous epimutations comes from work in *A. thaliana*, where the vast majority of methylation polymorphisms, particularly when considered as methylated regions, are shown to be stable across generations. However, some spontaneous changes in methylation loci (“epialleles”) become heritable, and are passed on to subsequent generations across meiosis (Hofmeister *et al.* 2017). The process of “epigenetic drift” may proceed alongside the process of genetic drift or differentiation while remaining unconnected mechanistically. A deeper understanding of the mutational processes as well as any non-clonal reproduction that may be occurring will be required to resolve the genetic and epigenetic contributions to invasion in this species.

## Acknowledgments

We thank Ramona Walls and Larry Gottschamer for help with original sample collections, Stony Brook University greenhouse staff Mike Axelrod and John Clumpp for maintaining the plants in the greenhouse, and Amy Litt at the New York Botanical Gardens for helping with new collections for the transcriptome sequencing. This work was supported by funding from the National Science Foundation (U.S.A.) DEB-1419960 and IOS- 1556820 (to CLR) and through the Global Invasions Network Research Exchange (Grant No. 0541673 for MR), Deutscher Akademische Austauschdienst (DAAD; MOPGA Project ID 306055 to CLR) and the Netherlands Organization for Scientific Research (NWO-ALW No. 820.01.025 to KJFV)

## Author Contributions Statement

CLR & KJFV conceived the study. CLR, KJFV, MA, and MR designed the experiments and analyses. MA, MR, CAMW and TVG did the epiGBS laboratory work. MA, MR, and TVG analyzed the epiGBS data. FY, DMA, WGF prepared, sequenced, and annotated the transcriptome. CLR, MA and MR wrote the first draft of the manuscript. All co-authors provided input and revisions to the manuscript.

## Data accessibility

R scripts and the specific version of the pipeline scripts we used for this study are available at: https://github.com/AlvarezMF/2020_Knotweed_epiGBS. Raw sequence data files for epiGBS (short reads after demultiplexing) have been, and PacBio (long reads) will be submitted to Dryad (https://doi.org/10.5061/dryad.zkh18938g) and NCBI. Links to this data will be made available at https://github.com/AlvarezMF/2020_Knotweed_epiGBS.

---

**Figure S1:**
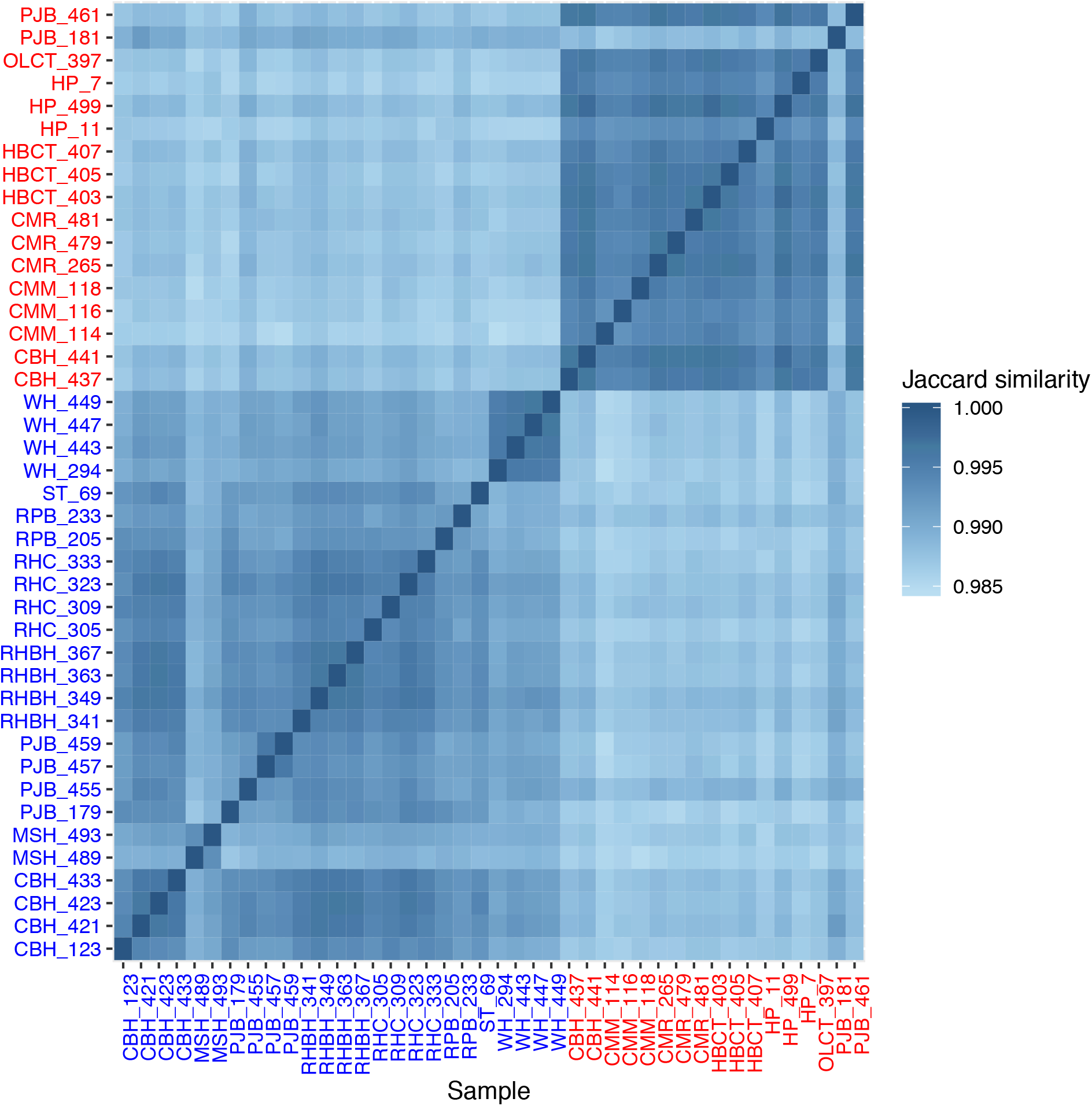
Heatmap of pairwise Jaccard similarities between individuals. *R. japonica* individuals are shown in blue, while *R. x bohemica* are shown in red. Sampling sites are identified by the first three letters of the sample names.

**Figure S2:**
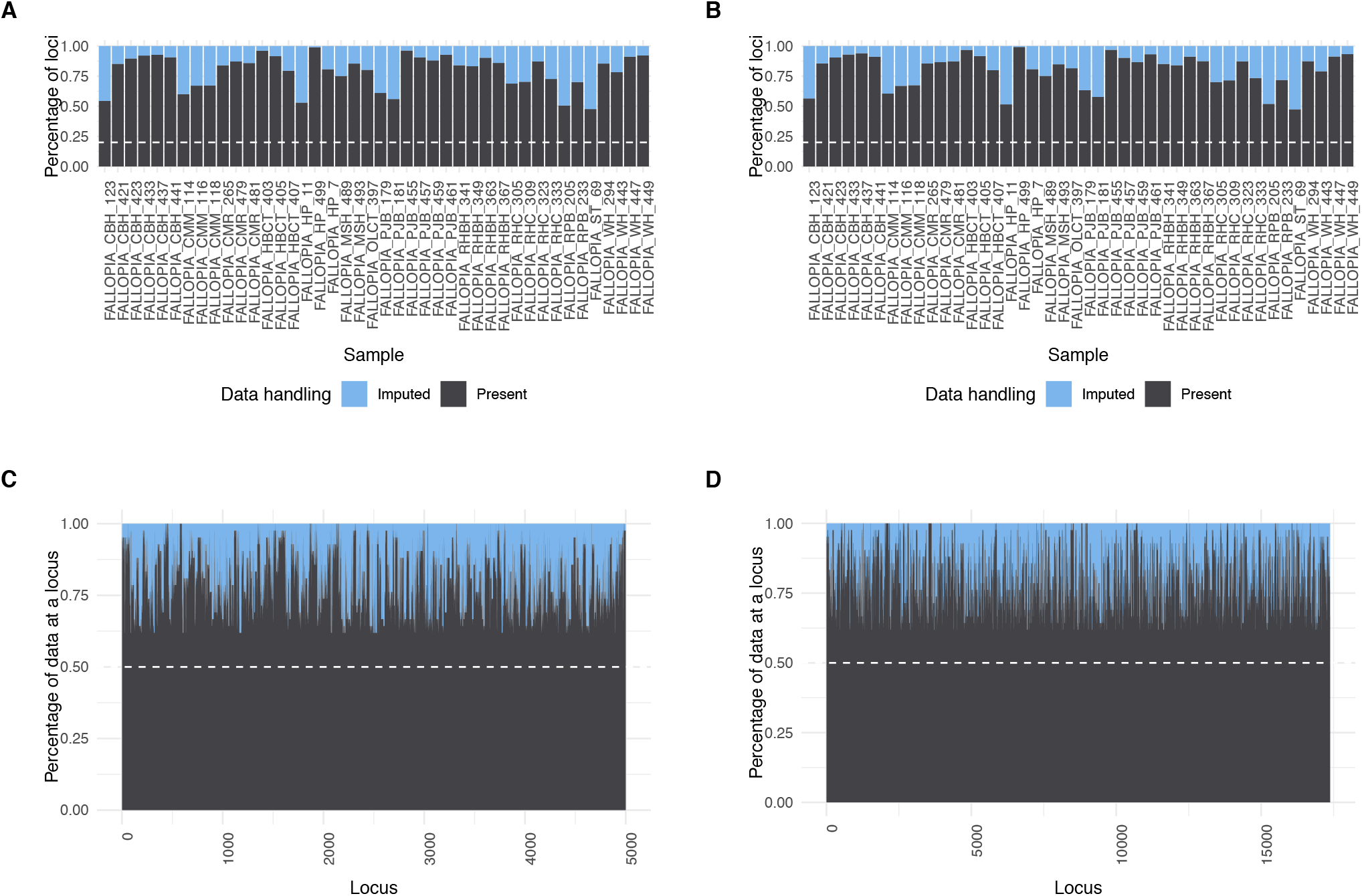
Amount of missing data imputed (light blue) and available (grey) by sample and by locus. Note that the locus plot shows 5000 loci drawn randomly from the group, and is shown as a representation of the general amount of missing data per locus.

## Notes

### Competing Interest Statement

The authors have declared no competing interest.

